# Adaptive Bayesian Optimization for State-Dependent Brain Stimulation

**DOI:** 10.1101/2023.04.30.538853

**Authors:** Sina Dabiri, Eric R. Cole, Robert E. Gross

## Abstract

Brain stimulation has become an important treatment option for a variety of neurological and psychiatric diseases. A key challenge in improving brain stimulation is selecting the optimal set of stimulation parameters for each patient, as parameter spaces are too large for brute-force search and their induced effects can exhibit complex subject-specific behavior. To achieve greatest effectiveness, stimulation parameters may additionally need to be adjusted based on an underlying neural state, which may be unknown, unmeasurable, or challenging to quantify *a priori*. In this study, we first develop a simulation of a state-dependent brain stimulation experiment using rodent optogenetic stimulation data. We then use this simulation to demonstrate and evaluate two implementations of an adaptive Bayesian optimization algorithm that can model a dynamically changing response to stimulation parameters without requiring knowledge of the underlying neural state. We show that, while standard Bayesian optimization converges and overfits to a single optimal set of stimulation parameters, adaptive Bayesian optimization can continue to update and explore as the neural state is changing and can provide more accurate optimal parameter estimation when the optimal stimulation parameters shift. These results suggest that learning algorithms such as adaptive Bayesian optimization can successfully find optimal state-dependent stimulation parameters, even when brain sensing and decoding technologies are insufficient to track the relevant neural state.

## I. Introduction

Brain stimulation and neuromodulation have recently emerged as an adjunct non-pharmacological treatment for neurological diseases such as Parkinson’s disease [1], depression [2] and epilepsy [3]. The effects of brain stimulation depend on many factors such as brain state [4], and the task the subject is performing [5]. In addition to the global brain state, methods that consider the local neural state could improve the efficacy and robustness of brain stimulation [6]. However, it is unknown how best to dynamically tailor stimulation to changes in neural state to improve the therapeutic effects of stimulation. Novel machine learning methods may be able to fill this gap.

An additional key challenge is that the stimulation parameter space (frequency, amplitude, pulse width, and contact location) is often too large for simple grid-based search strategies to be tractable [6]. Furthermore, each of combinations of these parameters can exhibit nonlinear interaction effects with different neural state variables, making state-dependent brain stimulation a challenging problem that requires efficient use of data.

One engineering approach that can improve upon grid search is Bayesian Optimization (BaO), where the maximum of an expensive cost function is found via setting a prior over the set of possible objective functions (optimum stimulation parameters), while the prior uses both exploration and exploitation. One of the ways to set the prior over the set of possible objective functions is to use a Gaussian Process (GP) model. Bayesian optimization then uses this prior to actively search for the input parameters that maximize the cost function.

Prior work from our laboratory has shown that neural state-dependent BaO (SDBO) can improve upon standard Bayesian optimization in finding state-dependent optimal stimulation parameters for brain stimulation when the neural state is known [6]. In addition, we have shown that using dimensionality reduction to reduce the neural-state can provide a condensed state-input for state-dependent BaO, allowing it to learn a complex or unknown relationship between the neural state and optimal stimulation parameters [7]. The dimensionality reduction method used was the t-distributed stochastic neighbor embedding (t-SNE) and the neural-state is the power spectral density. In further work we have shown that using a GP model for BaO reduces the number of samples needed to find optimal stimulation parameters by 33% and is appropriate for handling multi-objective reward functions in brain stimulation [8,9].

However, there are cases where the state cannot be observed or measured, but may still affect the brain’s response to stimulation. For example, the brain state may be distributed across many brain regions and cannot be measured with current chronically implanted devices, or cannot yet be reliably decoded from brain recordings. In this study, we investigate whether Bayesian optimization can be used to optimize non-stationary brain stimulation responses when the brain state is unknown, by incorporating an adaptive modeling strategy that prioritizes recent data over old data.

We use data from a rat optogenetic stimulation model to construct a simulation of a brain stimulation experiment where the stimulation response changes over time. We then use this simulation to compare how two implementations of adaptive Bayesian optimization and standard Bayesian optimization search for optimal stimulation parameters when given a non-stationary stimulation response.

## II. Methods

### A. Experimental Design and Data Collection

We use previously published data from an investigation of medial septum optogenetic stimulation and hippocampal electrophysiological recording in rats [6, 11]. The data was collected while a rat transitioned from anesthesia (2-3% isoflurane) to awake behavior. Stimulation parameters were applied in randomized order according to a grid of six frequencies (5, 7, 11, 17, 23, 29, and 35 Hz) and six amplitudes (0, 10, 20, 30, 40, and 50 mW/mm^2). The stimulation pulse waveform was a square wave delivered for 4 seconds, with a 4 second washout period between each trial.

### B. Bayesian Optimization

One of the components of Bayesian optimization is a Gaussian Process regression model, where the model optimizes stimulation parameters (pulse frequency [Hz] and optical amplitude [mW/mm2]) to achieve the ideal desired output, max (gamma power) given a neural-state measure (gamma power) [12]. All models are trained using the Python-based GPy package [13]. The Gaussian Process can approximate the objective function with a mean (m(x)) and covariance (k(x,x ’), also known as the kernel):

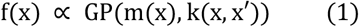

The kernel provides a weighting factor based on similarity between data points, such that points close to each other have greater influence over the model prediction than points that are farther apart. The kernel we used here is the Matern 5/2 kernel:

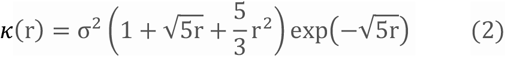

The function that does the sampling for next sample point based on the prior is the acquisition function. The acquisition function choses the next sample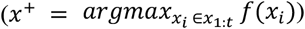, such that it would have the highest probability of improvement while having a trade-off parameter that increases the emphasis on exploration over exploitation as the number of samples increases. We use the upper-confidence bound acquisition function, where the next sample is chosen to maximize the mean of the Gaussian process model, plus the model’s standard deviation multiplied by a weighting factor:

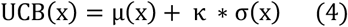

Furthermore, our implementation of the UCB acquisition function includes an instantaneous regret function 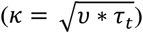in the acquisition function weighting. Experimentally the parameter *v* is set at 0.1, and 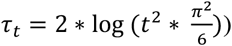.

### C. Adaptive Modeling for Bayesian Optimization

We evaluated two implementations of an adaptive Bayesian optimization approach:

Dynamic BaO: A forgetting factor is incorporated into the implementation of BaO, such that only the 20 most recent samples are used to fit the GP at each iteration.

Adaptive BaO: Time is directly incorporated as a third input to the GP kernel used in BaO, in addition to stimulation parameters. At each iteration, the current iteration number is used to condition the Gaussian process, and the acquisition function is applied only to the remaining two-dimensional model GP cross-section. This method uses the kernel to reweight the importance of all prior data points based on their temporal distance from the current time step.

### C. Simulation Model

To design a state-dependent simulation for this study, we extend our lab’s previously published framework for designing optimization algorithms *in silico* based on collected experimental data [9-10]. A GP model was trained to predict hippocampal gamma power (32-50 Hz; summed across recording channels) during stimulation, as a function of three variables: stimulation frequency, amplitude, and the pre-stimulation gamma (32-50 Hz; also summed across recording channels) neural state. As shown in our previous work, stimulation in this system has state-dependent properties: as the brain state changes from anesthetized to awake, the optimal stimulation parameters that maximize gamma power shift from maximum amplitude and gamma-range (40 Hz) frequency to maximum amplitude and half-gamma (∼20 Hz) frequency. This change can be predicted by pre-stimulation gamma power, which is correlated with the animal’s anesthesia level. This shift in the optimal stimulation frequency will be used as the ground truth neural state change to evaluate how well each algorithmic method of this study can adapt to a non-stationary stimulation response.

The pre-stimulation gamma neural state data was then used to construct a manual time series input that provides the changing state of the simulation (Figure 1a). Over 100 iterations, the state is linearly adjusted from minimum gamma power (n = 0) to its maximum value (n = 50), then back to its minimum value (n=100), indicating a shift of awake ⇒ anesthesia ⇒ awake. The simulation model can also be sampled at any given input of stimulation parameters and neural state to provide a noisy, normally distributed response that reflects the statistical distribution of values observed in the original experiment:

**Figure 1.**
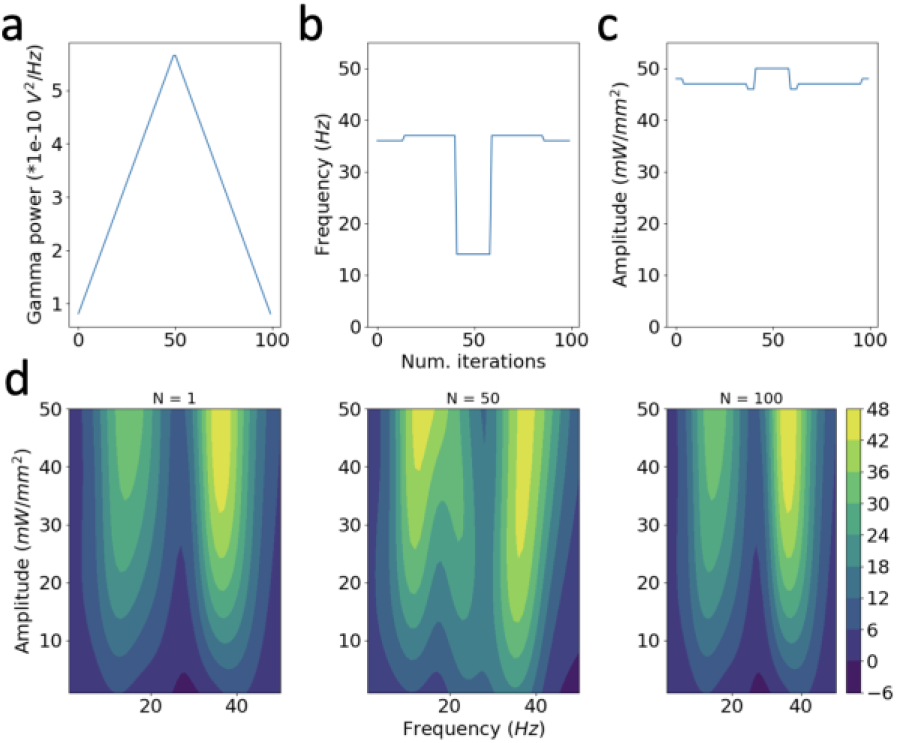
Visualization of the state-dependent stimulation model. a) Time series of the manual state input applied to the model vs. 100 simulation iterations. b-c) Ground-truth stimulation frequency (b) and amplitude (c) that maximize the simulation model’s mean prediction.d) Surface plot of the simulation model’s mean prediction vs. stimulation parameters at different time steps of the simulation, corresponding to minimum (1 iteration, left), maximum (50 iterations, center), and minimum (100 iterations, right) neural state values.

Over the course of the 100-sample simulation, Bayesian optimization is used to select stimulation parameters for sampling, and the neural state time series provides the state input. At each iteration, a gamma power output is sampled from the model at the given combination of inputs and provides the gamma power sample from the objective function that is used for both BaO algorithms. Furthermore, the simulation model will serve as the ground truth for the Static and Dynamic model to compare to (Figure 1 right panel).

### E. Simulation Design

All three BaO methods are trained without the pre-stimulation neural state, and are only provided the frequency, amplitude, and gamma power output sampled from the simulation model. Each model is first trained using 20 burn-in points where the stimulation parameters are randomly selected, followed by the UCB acquisition function determining sampling throughout the remainder of the simulation using a grid of frequencies from 1-42 Hz and amplitudes 1-50. The simulation is 100 iterations long and repeated for 30 trials per algorithm.

## III. Results

## A. Description of the State-Dependent Simulation Model

Optimal parameters and mean surface of the simulation Gaussian process model are visualized in Figure 1b-d. The gamma power response is characterized by the greatest increase at stimulation parameters of maximum amplitude and frequency. A smaller peak at the subharmonic frequency of the maximum is also present in the model at awake states and increases during anesthesia (iterations 40-60). In this period, the frequency that maximizes the gamma power output of the simulation model transitions to <20 Hz, whereas the optimal amplitude remains similar.

### B.Performance of Static and Adaptive Search Strategies

Figure 2 demonstrates the mean parameters chosen by the acquisition function for sampling by each BaO method (yellow) and the parameters each method estimates will maximize the simulation model’s output (blue) during each iteration of the simulation. Static BaO quickly converges onto the optimum of the simulation and has a small standard deviation of the parameters chosen by the acquisition function, indicating high exploitation and low exploration. After the state change at 40 iterations, both adaptive methods display a higher standard deviation of sampled parameter values, indicating an increase in exploration. The adaptive methods find the optimal frequency of 40 Hz in the beginning of the simulation, and additionally decrease the optimal frequency estimate when the optimal frequency shifts at 40 iterations.

**Figure 2.**
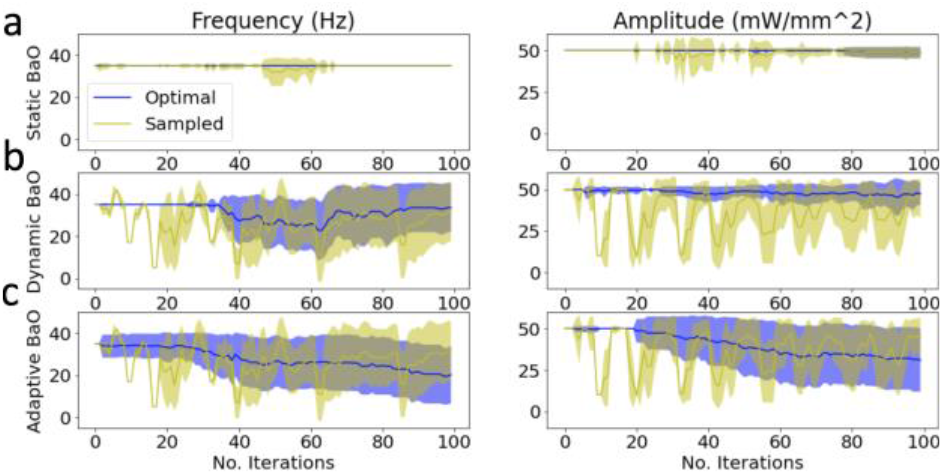
Static, dynamic and adaptive Bayesian optimization simulation performance. A) Time series of frequency (left) and amplitude (right) parameters modeled by Static BaO throughout the simulation duration. Blue: parameter value that the BaO Gaussian process regression model estimates will produce the maximum gamma power response. Yellow: parameter value that the acquisition function selected for sampling. Shading indicates mean +/-standard deviation across 30 repeated runs of the simulation. B-C) Equivalent time series for parameters modeled by dynamic (B) and adaptive (C) Bayesian optimization.

### C. Adaptive Bayesian Optimization Outperforms Static Optimization in Non-stationary Search

Figure 3a quantifies the standard deviation of both frequency and amplitude values sampled by each method, showing that dynamic and adaptive BaO outperform static BaO in exploration. Both adaptive methods identify an optimal frequency that is closer to the simulation model’s optimal frequency during the state change. Dynamic BaO more consistently finds the optimal frequency at the beginning and end of the simulation than Adaptive BaO.

**Figure 3.**
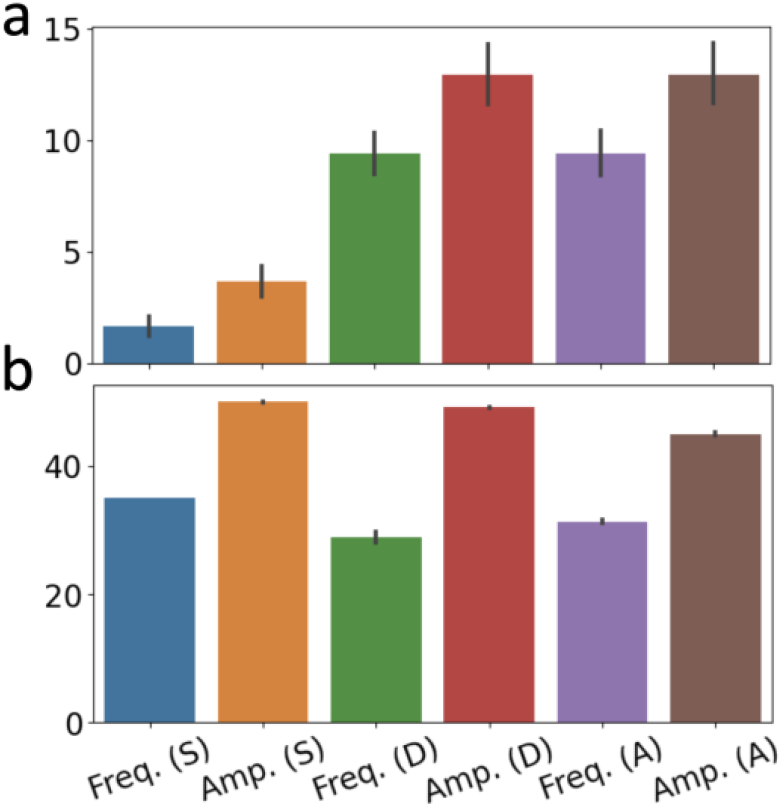
Results of parameters chosen by static and adaptive Bayesian optimization. a) Standard deviation of parameters chosen for sampling by static (S), dynamic (D) and adaptive (A) BaO methods. b) Mean value of estimated optimal parameters after iteration 40 of the simulation for each method. All bars indicate mean +/-95% confidence interval.

## IV. Discussion

In this study, we used rat optogenetic stimulation data to construct a simulation of a state-dependent brain stimulation experiment and used it to characterize the performance of both static and adaptive Bayesian optimization. We showed that incorporating adaptation into Bayesian optimization can allow it to better learn state-dependent changes of optimal parameters in the brain stimulation response.

One limitation in this study is the assumption of a Gaussian process for modeling the experimental data, which in a real experiment may have more complex and nonlinear dynamics. The two computational mechanisms used for adaptation in this study also have both advantages and limitations; the computational simplicity of a forgetting factor means that this implementation could be suitable for use with implanted devices. The kernelized method of Adaptive BaO may allow for more data-efficient inference by incorporating all data, but could suffer from computational speed issues as fitting methods for Gaussian process regression models scale with cubic time complexity vs. the size of the data set used [13]. Interestingly, this method did not clearly outperform the more simple forgetting factor, as it may be influenced by “memory” of old data that constrains flexibility when encountering new data. Additionally, the forgetting factor and length scale values for adaptation were selected *a priori* and would need to be selected for real-time applications based on an expected timescale of state change in the system or estimated from prior data before deployment. In addition, constant adaptation hyperparameters may be insufficient if the speed at which the neural state changes can vary over time.

This latter limitation could be overcome by incorporating a Kalman filter-based method to estimate a latent state variable for the neural state in real-time, or by using other methods to empirically adjust the value of the forgetting factor throughout the experiment based on measured data [15]. The choice of Bayesian optimization for the basis of the adaptive algorithm could also be supplanted or improved by more recent advancements in machine learning. Reinforcement learning algorithms may be less sample-efficient than Bayesian optimization in finding a maximizing set of input parameters, but they may be suitable for online applications; sample-efficiency and other gaps could be improved upon by using recent advancements in meta-learning for active and reinforcement learning [16-17].

Multiple different types of non-stationarities and state-dependent properties observed in different types of neural systems could require different machine learning strategies for adaptation to achieve best performance. Common sources of non-stationarities may include physical movement or damage to recording electrodes, buildup of scar tissue around implanted or stimulated sites, behavioral or task-related changes in brain state, and dynamic neural changes induced by a stimulation pulse train such as synaptic exhaustion [4, 18-20]. Many such applications have been addressed with Kalman-inspired tracking methods and are characterized by slow and sustained rather than rapid or frequently changing dynamics, suggesting that methods assuming conserved change could be suitable for a wide array of applications in state-dependent brain stimulation [18].

